# Recurrent connections between CA2 pyramidal cells

**DOI:** 10.1101/321513

**Authors:** Kazuki Okamoto, Yuji Ikegaya

## Abstract

Recurrent excitatory synapses are shown theoretically to play roles in memory storage and associative learning, and such recurrent synapses are well described to occur in the CA3 region of the hippocampus. Here, we report that the CA2 region also contains recurrent excitatory monosynaptic couplings. Using dual whole-cell patch-clamp recordings from CA2 pyramidal cells in mouse hippocampal slices under differential interference contrast microscopic controls, we evaluated monosynaptic excitatory connections. Unitary excitatory postsynaptic potentials occurred in 1.4% of 502 cell pairs. These connected pairs were located preferentially in the superficial layer and proximal part (CA2b) of the CA2 region. These results indicate that recurrent excitatory circuits are dense in the CA2 region as well as in the CA3 region.

## Introduction

Theoretical studies have suggested that reentrant positive-feedback excitation of the neuronal network is critical for nonlinear information processing, including memory storage, associative learning, and pattern separation/completion (Amari et al., 1996; Hebb, 1949; Hopfield, 1982; Kohonen, 1998). The hippocampal CA3 region is one of the candidate brain regions that anatomically contain self-associative excitatory networks and functionally exert such information processing (Guzman et al., 2016; MacVicar and Dudek, 1981; Miles and Wong, 1986; Nakazawa et al., 2004; Treves and Rolls, 1994). However, only a few studies have directly measured monosynaptic recurrent connections in the hippocampus (Deuchars and Thomson, 1996; Guzman et al., 2016; Miles and Wong, 1986), particularly in the CA2 region (Mercer et al., 2012).

The CA2 region of the hippocampus is unique in terms of spatial representation and social memory (Hitti and Siegelbaum, 2014; Mankin et al., 2015), but it has been less investigated because it is not included in the classical tri-synaptic pathway and because the CA2 region is anatomically small, which limits experimental access. Notably, whether CA2 pyramidal cells are mutually excited at the monosynaptic level remains controversial. Some reports suggest that the CA2 region contains recurrent excitatory circuits (Lu et al., 2015) because CA2 neurons elaborate highly arborized dendrites (Dudek et al., 2016) and because CA2 neurons can initiate sharp wave/ripple oscillations (Oliva et al., 2016).

In the present study, we directly measured excitatory monosynaptic connections between CA2 pyramidal cells using double whole-cell patch-clamp recordings. We observed monosynaptic unitary excitatory postsynaptic potentials (EPSPs) between 1.4% of the CA2 pyramidal cell pairs. The CA2-to-CA2 connections existed preferentially in the CA2b subarea and exhibited a preferential spatial direction.

## Results and Discussion

Recent histological studies have broadened the conventional definition (Lorente de Nó, 1934) of the CA2 region based on specific molecular expressions, such as those of G-protein signaling protein 14 (RGS14) and striatal-enriched protein tyrosine phosphatase (STEP) (Dudek et al., 2016; Kohara et al., 2014; Lein et al., 2005; Noguchi et al., 2017). We also confirmed that RGS14 immunoreactivity indicated the macroscopic location of the CA2 region in transverse slices of the hippocampus. At the single cell level, however, RGS14-positive cells were sparse near the CA1/CA2 and CA2/CA3 borders (Figure 1). This sparsity was found particularly in the ventral hippocampus. This salt-and-pepper distribution of RGS14-positive cells indicates that a significant number of non-CA2 pyramidal cells (presumably, CA1 or CA3 pyramidal cells) are present in the CA2 region.

**Figure 1.**
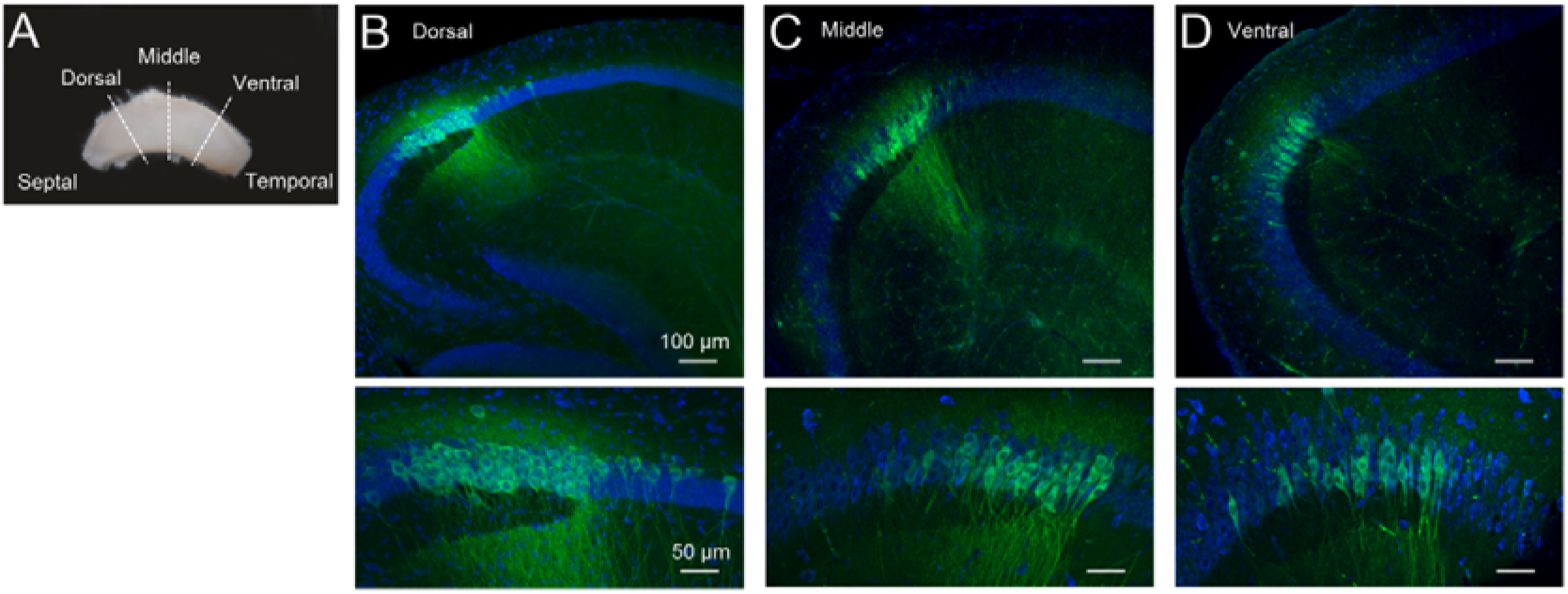
Distributions of CA2 pyramidal cells along the longitudinal axis of the mouse hippocampus. **A)** A whole hippocampus was sectioned transversely at the dorsal, middle, and ventral areas, indicated by the broken white lines. **B-D)** The transverse sections at the dorsal (B), middle (C), and ventral (D) areas were labeled with an anti-RGS14 antibody (*green*, a CA2 marker) and NeuroTrace 435/455 Nissl Stain (*blue*). The bottom photos are magnified images of the CA2 parts, indicating that the CA2 borders next to the CA1 and CA3 regions are ambiguous because of the sparse distribution of the CA2 pyramidal cells, particularly in the ventral hippocampus.

We investigated recurrent CA2 connectivity by recording unitary EPSPs between two CA2 pyramidal cells. We visually targeted pyramidal cells using a differential interference contrast microscope. The recorded neurons were intracellularly filled with biocytin through patch-clamp pipettes and were then immunostained using an antibody against RGS14 or STEP (Figure 2A). A train of four action potentials was evoked at 20 Hz in one of the recorded cells, and the evoked membrane potentials were monitored for the other cells (Figure 2B). In the 502 pairs tested, we found 7 chemical synaptic connections and no electrical coupling; that is, the CA2-to-CA2 synaptic connection probability was 1.4%. The unitary EPSPs had a mean peak amplitude of 0.43 ± 0.14 mV and a mean transmission failure rate of 54 ± 24% (mean ± SD of 7 connections). Three of the 7 connections exhibited short-term depression in response to four presynaptic spikes (Figure 3). We also patched 108 CA3 pyramidal cell pairs and found 3 chemical synaptic connections (2.8%), whose unitary EPSPs had a mean peak amplitude of 0.75 ± 0.54 mV and a mean failure rate of 45 ± 22% (3 connections). These parameters of the CA3 pairs are similar to those reported in previous studies of CA3 connections (Guzman et al., 2016; Miles and Wong, 1986), and they did not differ significantly from those of the CA2 connections in the present study (*P* > 0.05, Student’s *t*–test).

**Figure 2.**
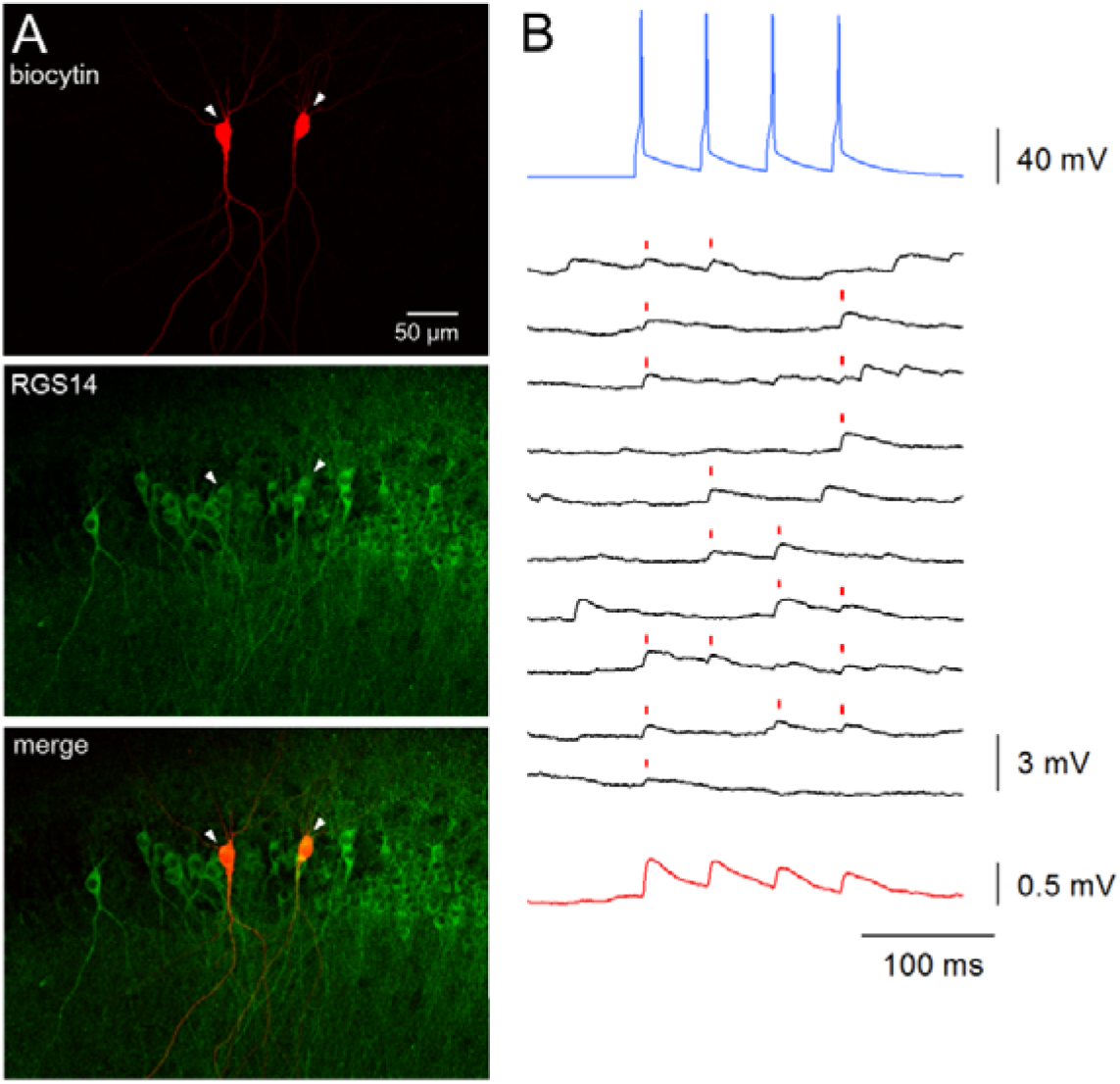
Monosynaptic recurrent connection of a CA2-CA2 pair. **A)** *Top* : Confocal images of a synaptically connected pair of CA2 pyramidal cells that were filled with biocytin (*red*). *Middle* : The CA2 region was immunostained for RGS14 (*green*). *Bottom* : RGS14 was expressed in the recorded cells. **B)** Unitary EPSPs of the connected pair. *Top* : Mean trace of a train of current injection-induced action potentials of the presynaptic cell (blue, averaged for 50 traces). *Middle* : Representative postsynaptic membrane responses recorded in the current-clamp mode for the successive trials (black). *Bottom* : an average of 50 recorded traces for the postsynaptic cell.

**Figure 3.**
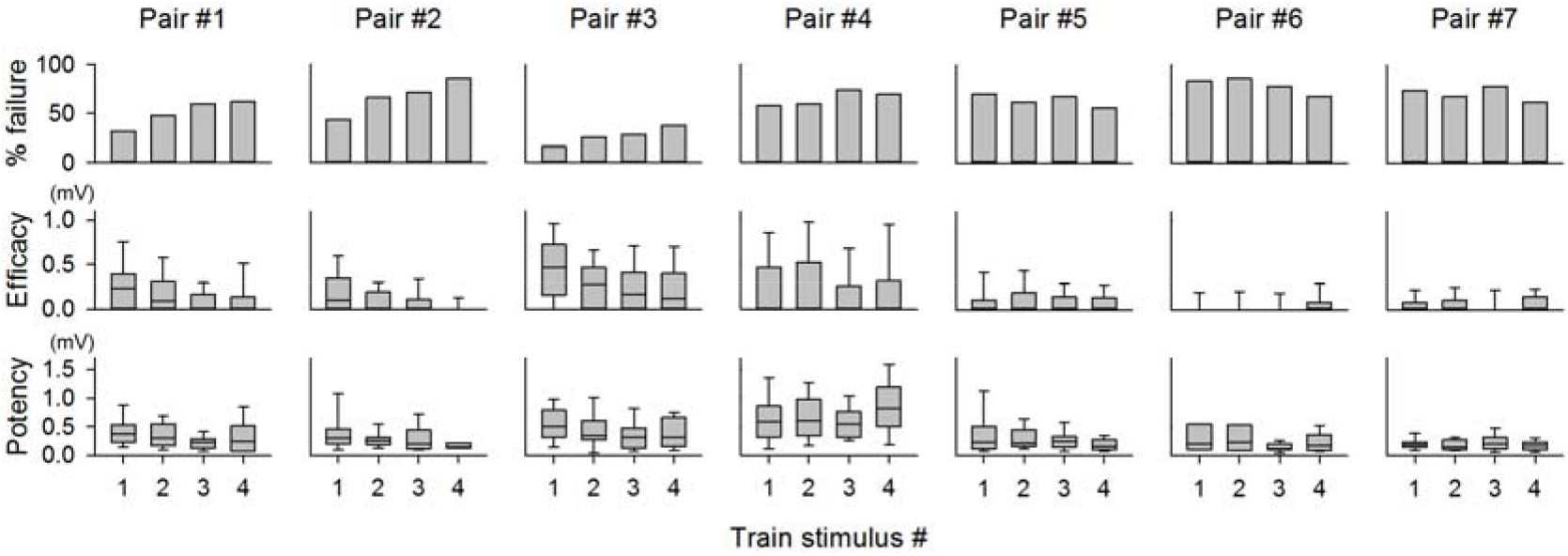
Unitary EPSP properties of all connected pairs. *Top:* Percentages of synaptic failures in the responses to a presynaptic spike train (4 pulses at 20 Hz) for 7 connected pairs. Train stimuli were repeated 50 times. *Center:* Synaptic efficacies across the 50 trains (including failure events). *Bottom:* Synaptic potencies (without failure events).

During the whole-cell recordings, we measured the physical distances between the somata of two recorded cells in the images taken with a differential interference contrast microscope. The inter-soma distances were not correlated with the connection probability (Figure 4). This result appears similar to the previously reported spatial patterns of CA3 connectivity (Guzman et al., 2016) but is different from those of neocortical connectivity, in which more adjacent pairs had a higher probability of connections (Peng et al., 2017; Perin et al., 2011; Song et al., 2005). When considering the connectivity, however, one needs to consider the anatomical size of the CA2 region; the majority of CA2 cell pairs have inter-soma distances of less than 100 µm; thus, we cannot rule out the possibility that this sampling restriction masked the distance dependence.

**Figure 4.**
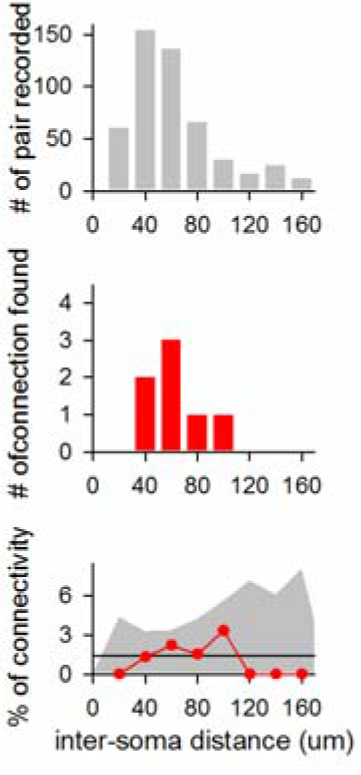
Lack of correlation of CA2-CA2 connections with their inter-cell distances. The top histogram indicates the distributions of the distances between the centroids of the cell bodies of all recorded pairs. The center histogram indicates the same distribution as the top, but for only the synaptically connected pairs. The bottom graph plots the connection probabilities (*i.e*., the center histogram divided by the top histogram) as a function of the inter-soma distance (red). The black line is the mean connection probability (1.4%). The gray area is the 95% confidence interval estimated by 10,000 randomly resampled surrogates.

CA1 and CA3 pyramidal cells are heterogeneous along both transverse (Ishizuka et al., 1990; Lee et al., 2015; Lu et al., 2015; Sun et al., 2017) and radial axes (Kohara et al., 2014; Lee et al., 2014; Valero et al., 2015). Similar heterogeneity has also been reported in the CA2 region (Oliva et al., 2016). We thus investigated whether CA2-to-CA2 connections are spatially biased in the CA2 stratum pyramidale. For one of the 7 connections, we failed to accurately identify the relative loci of the recorded cells in the RGS14-positive CA2 region. Thus, we analyzed the remaining 6 pairs. The relative location of each recorded cell was determined in the CA2 stratum pyramidale, which was rectangularly standardized according to the RGS14-positive or STEP-positive areas (Figure 5A). In this cell map, we noticed two structural tendencies. First, the directions from presynaptic cells to postsynaptic cells were spatially biased (Figure 5B; *V* = 2.05, *P* = 0.02, *n* = 6 pairs, *V*-test). A given CA2 pyramidal cell tended to make a synaptic connection with a CA2 pyramidal cell that was more proximal to the CA3 regions. This tendency suggests that compared to distal CA2 cells, more proximal CA2 pyramidal cells receive more recurrent inputs from other CA2 pyramidal cells.

**Figure 5.**
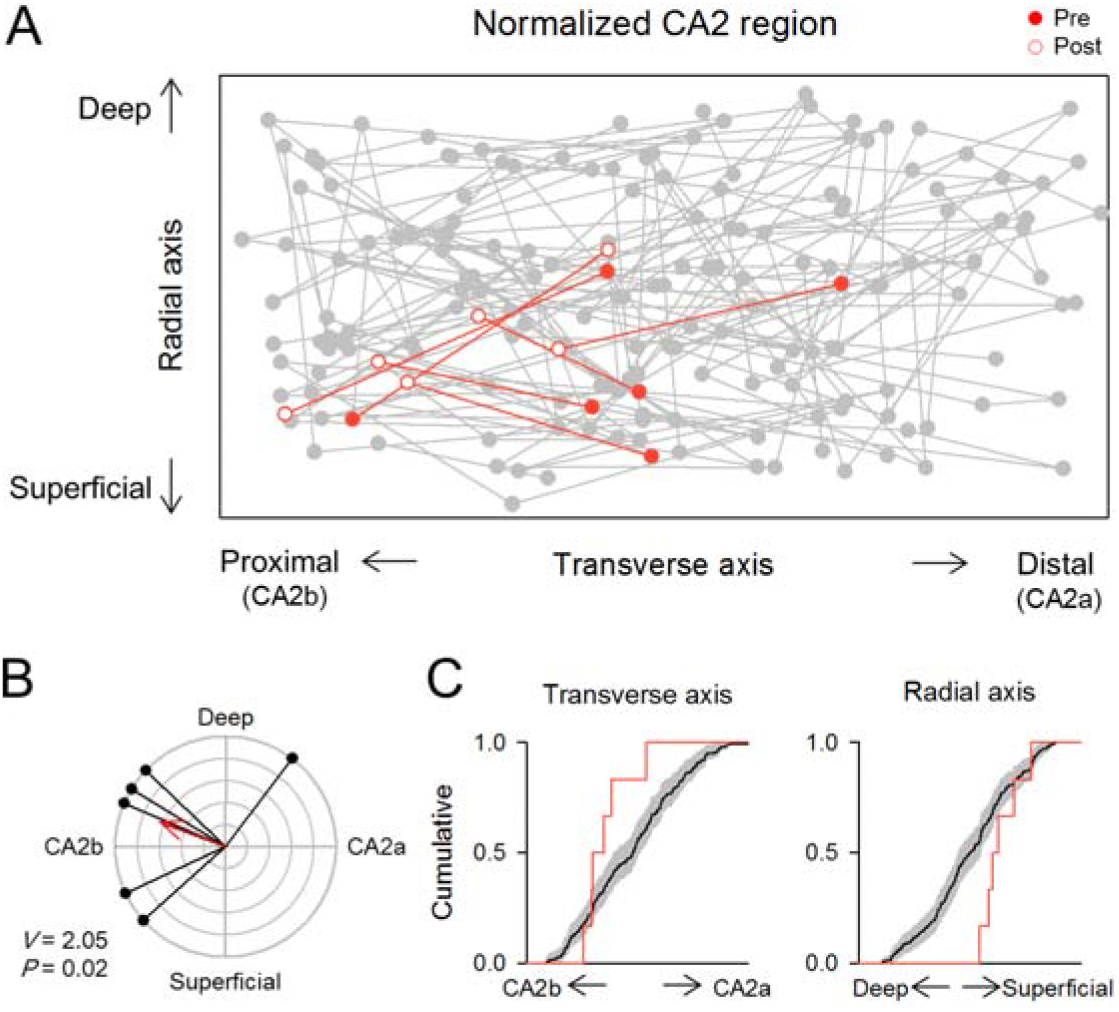
Spatial bias of CA2-CA2 connections. **A)** The stratum pyramidale of the CA2 region was normalized as a rectangle according to RGS14-positive or STEP-positive areas. Gray dots and lines indicate the soma location of the recorded CA2 pyramidal cells and their potential connections (unconnected), respectively. Red lines represent the synaptic connections (n = 6). Red filled and open circles indicate the soma location of the presynaptic and postsynaptic cells, respectively. **B)** A polar plot of the inter-soma directions of the synaptically connected pairs. Each black line indicates the direction from the soma of one presynaptic cell to the soma of its postsynaptic cell. The red arrow represents the mean vector of all 6 connected pairs. *V* = 2.05, *P* = 0.02, *V*-test. **C)** Cumulative distributions of the midpoints between two soma locations of connected pairs (*red line*) and unconnected pairs (*black line*) along the transverse axis (*left*) and the radial axis (*right*). The gray areas indicate the 95% confidence intervals estimated by Kaplan-Meier method.

Second, the connected pairs were located preferentially in the superficial layer and the proximal side of the CA2 region (Figure 5C). The difference between the superficial and deep layers of the stratum pyramidale has not been well studied in the CA2 region. A recent study demonstrated that the CA2 deep layer initiates sharp wave/ripple oscillations (Oliva et al., 2016). Recurrent excitation is proposed as one of the substrates of sharp wave/ripple generation (Traub et al., 1989), but our results indicate that the recurrent connections of the CA2 pyramidal cells are nearly exclusively located in the superficial layer. The generation of sharp waves/ripples is a complex process, including dense recurrent circuits, timed interneuron activation (Stark et al., 2014) and extrahippocampal controls (Buzsaki, 2015). The CA2 region may trigger sharp waves/ripples independently of local recurrent excitation.

We found that the proximal CA2 (CA2b) subarea, but not the distal CA2 (CA2a) subarea, has dense recurrent connections, similar to the CA3 region. According to the most recent definition (Dudek et al., 2016; Lein et al., 2005), the CA2b subarea encompasses the stratum lucidum and receives monosynaptic inputs from the dentate gyrus (Kohara et al., 2014; Sun et al., 2017); however, the unitary EPSP sizes are smaller in CA2 pyramidal cells than in CA3 pyramidal cells (Sun et al., 2017). In addition, CA2 pyramidal cells receive strong excitatory inputs from layer III neurons in the entorhinal cortex (Chevaleyre and Siegelbaum, 2010). These extrahippocampal inputs contribute to the unstable dynamics of CA2 neuronal activity (Lee et al., 2015; Lu et al., 2015; Mankin et al., 2015). The CA3 network contains clustered circuit motifs (Guzman et al., 2016). The neocortex also contains rich clustered connections (Peng et al., 2017; Perin et al., 2011; Song et al., 2005). In CA2 pyramidal cells, we did not find bidirectionally connected pairs. This may be because the numbers of pair recordings are simply not enough.

A previous study reported a very low connection probability (0.22%) between CA2 pyramidal cells (Mercer et al., 2012); however, this study did not identify CA2 pyramidal cells histologically using the CA2 cell markers (RGS-14 or STEP). Given that CA2 pyramidal cells are sparse even in the CA2 pyramidal cell layer, the authors may have underestimated the true CA2 connectivity. Another possibility is that they might have focused more on the CA2a subarea, which was conventionally defined as “the CA2 region”. This sampling bias could also lead to an underestimation of the CA2 connectivity.

The CA2 region was reported to contain a higher density of GABAergic interneurons than the CA1 or CA3 region (Leranth and Ribak, 1991; Mercer et al., 2007). Interestingly, within the CA3 region, the CA3a subarea receives more inhibitory inputs and has denser recurrent connectivity than the CA3b subarea (Sun et al., 2017).

These observations suggest similar circuit properties of the CA2b and CA3a subareas. The high density of recurrent excitatory connections may be counter-balanced by strong inhibitory inputs and maintain the excitatory-to-inhibitory balance in the CA2 local circuit.

## Methods

### Animal experiment ethics

Experiments were performed with the approval of the Animal Experiment Ethics Committee at the University of Tokyo (approval no. P29–9) and according to the University of Tokyo guidelines for the care and use of laboratory animals. Institute of Cancer Research (ICR) mice (SLC) were housed in cages under standard laboratory conditions (12 h light/dark cycle, ad libitum access to food and water). All efforts were made to minimize the animals’ suffering and the number of animals used.

### Acute slice preparation

Acute slices were prepared from the hippocampi of ICR mice (17–26 postnatal days). The mice were anesthetized with isoflurane and then decapitated. The brains were removed and placed in an ice-cold oxygenated solution consisting of (in mM) 222.1 sucrose, 27 NaHCO_3_, 1.4 NaH_2_PO_4_, 2.5 KCl, 1 CaCl_2_, 7 MgSO_4_, and 0.5 ascorbic acid. The brains were sliced horizontally at a thickness of 400 µm using a VT1200S vibratome (Leica). The slices were allowed to equilibrate at room temperature for at least 0.5 h while submerged in a chamber filled with oxygenated aCSF consisting of (in mM) 127 NaCl, 26 NaHCO_3_, 1.6 KCl, 1.24 KH_2_PO_4_, 1.3 MgSO_4_, 2.4 CaCl_2_, and 10 glucose. The slices were mounted in a recording chamber and perfused at a rate of 1.5–3 ml/min with oxygenated aCSF.

### In vitro electrophysiology

All recordings were performed at 32−34°C. Whole-cell recordings were collected from up to four CA2 pyramidal cells using a MultiClamp 700B amplifier and a Digidata 1550 digitizer controlled by pCLAMP10.6 software (Molecular Devices). Borosilicate glass pipettes (3−6 M Ω) were filled with a solution containing (in mM) 135 K-gluconate, 4 KCl_2_, 0.3 EGTA, 10 HEPES, 10 Na_2_-phosphocreatine, 4 MgATP, 0.3 Na_2_GTP and 2.0 biocytin. The signals were gained 10-fold, low-pass filtered at 1 kHz and digitized at 20 kHz. The existence of synaptic connectivity was assessed by averaging 50 successive traces in which 4 spikes at 20 Hz were induced by current injection in presynaptic cells.

### Histology

CA2 pyramidal cells were perfused intracellularly with 2 mM biocytin for whole-cell recordings. After the recordings, the slices were fixed for at least 20 h at 4°C in 0.1 M Na_3_PO_4_, pH 7.4, containing 3% (w/v) formaldehyde. The sections were incubated with 2 μg/mL streptavidin-Alexa Fluor 594 conjugate and 0.2% Triton X-100 for 6 h, followed by incubation with 0.4% NeuroTrace 435/455 blue-fluorescent Nissl Stain (Thermo Fisher Scientific; N21479) overnight. The tissue sections were incubated subsequently with mouse primary antibodies for RGS-14 (NeuroMab; N133/21; 1:500) or STEP (Cell Signaling Technology; 4396S; 1:500) for 16 h at 4°C, followed by incubation with a secondary goat antibody to mouse IgG (Thermo Fisher Scientific; A-11001; 1:500) for 6 h at 4°C.

## Acknowledgments

We thank Dr. Segundo Jose Guzman, Dr. Vivien Chevaleyre, Dr. Rebecca Piskorowski, and Dr. Takeshi Sakaba for their comments on the experiment. This work was supported by Grants-in-Aid for Scientific Research (17K19441 & 18H05114) and by the Human Frontier Science Program (RGP0019/2016). This work was conducted partly as a program at the International Research Center for Neurointelligence (WPI-IRCN) of the University of Tokyo Institutes for Advanced Study at the University of Tokyo.

